# Semi-supervised machine learning for sensitive open modification spectral library searching

**DOI:** 10.1101/2022.09.30.510366

**Authors:** Issar Arab, William E. Fondrie, Kris Laukens, Wout Bittremieux

**Affiliations:** Department of Computer Science, University of Antwerp, Antwerp, Belgium; Biomedical Informatics Network Antwerpen (biomina), Antwerp, Belgium; Talus Bioscience, Seattle, WA, USA; 98195, USA; Skaggs School of Pharmacy and Pharmaceutical Sciences, La Jolla, CA, USA

## Abstract

A key analysis task in mass spectrometry proteomics is matching the acquired tandem mass spectra to their originating peptides by sequence database searching or spectral library searching. Machine learning is an increasingly popular post-processing approach to maximize the number of confident spectrum identifications that can be obtained at a given false discovery rate threshold. Here, we have integrated semi-supervised machine learning in the ANN-SoLo tool, an efficient spectral library search engine that is optimized for open modification searching to identify peptides with any type of post-translational modification. We show that machine learning rescoring boosts the number of spectra that can be identified for both standard searching and open searching, and we provide insights into relevant spectrum characteristics harnessed by the machine learning model. The semi-supervised machine learning functionality has now been fully integrated into ANN-SoLo, which is available as open source under the permissive Apache 2.0 license on GitHub at https://github.com/bittremieux/ANN-SoLo.

## 1 Introduction

To identify peptides from tandem mass spectrometry (MS/MS) data, the experimental spectra are compared to theoretical spectra simulated from peptides derived from a sequence database^1^ or previously acquired reference spectra,^2,3^ after which the corresponding peptide labels are transferred to high-quality matches. To ensure that only correct peptide-spectrum matches (PSMs) are accepted, a target–decoy strategy^4^ is typically employed, during which the spectra are not only searched against peptides of interest, but also against decoy peptides that by definition cannot occur. The number of matches to decoy peptides at a specific score threshold can then be used as a proxy for the number of incorrect target matches, and only those matches at a controlled false discovery rate (FDR) are accepted.^5^

Originally introduced by Percolator,^6^ an increasingly popular approach to expand upon the target–decoy searching strategy is to use machine learning for PSM rescoring. Percolator uses a semi-supervised support vector machine (SVM) to discriminate between confident and decoy spectrum identifications, by iteratively learning patterns that correspond to confident PSMs. This makes it possible to improve the spectrum identification results by accepting more target PSMs at a specific FDR threshold. Since the original description of Percolator,^6^ several approaches that extend upon this idea have been proposed, including tools that use different machine learning models instead of a linear SVM,^7–9^ integration with various search engines,^10–12^ and additional features that the classifier can use, often powered by deep learning.^13–20^

Another recent development is the emergence of novel search engines that can efficiently perform so-called open modification searching (OMS).^21^ An open search uses a wide precursor mass tolerance, on the orders of hundreds of Dalton, to identify not only peptides with a matching precursor mass, but also peptides whose mass differs from the precursor mass due to the presence of post-translational modifications (PTMs).^22,23^ In this fashion, variable PTMs no longer need to be specified or known in advance, but instead can be derived from the observed precursor mass differences, for the unbiased identification of modified peptides. Although traditional OMS approaches necessarily restricted the search space to only a handful of proteins due to computational and runtime constraints,^24^ recently several new search engines have implemented efficient approaches to perform open searches.^25–31^

Here, we have integrated PSM rescoring into the ANN-SoLo spectral library search engine,^29,30^ a tool for efficient open modification searching. ANN-SoLo uses a cascade search strategy^32^ to optimally identify both unmodified and modified peptides: in the first stage a standard search is performed to identify unmodified peptides, after which the remaining unidentified spectra are submitted to the second stage during which an open search is performed to additionally identify modified peptides. We have augmented this approach by natively integrating PSM rescoring into ANN-SoLo using the mokapot Python framework for semi-supervised learning for peptide detection.^33^ We show that PSM rescoring increases the number of spectra that are identified by ANN-SoLo and boosts the spectrum identification sensivity, both for standard searching and open searching.

## 2 Methods

### 2.1 Integration of semi-supervised PSM rescoring in ANN-SoLo

PSM rescoring has been implemented in ANN-SoLo using the mokapot Python framework.^33^ Features for each PSM are derived from the match between a query MS/MS spectrum and a library MS/MS spectrum and include: the length of the peptide sequence, the precursor charge (one-hot encoded between two and five or more), the query/library spectrum precursor m/z, the (absolute) precursor *m/z* difference in parts per million, the (absolute) precursor *m/z* difference in Dalton, the cosine similarity,^29^* the number of matching fragment ions, the fraction of matching fragment ions for the query/library spectrum,* the fraction of covered intensity for the query/library spectrum,* the mean squared *m/z* error of the matching fragment ions,* the mean squared intensity error of the matching fragment ions,* the spectral contrast angle,^34^* the hypergeometric score of obtaining more than the observed number of matching fragment ions by random chance,^35^ the Kendall-Tau score of correspondence between the intensity ranks of the matching fragment ions,^35^ the MSforIDv1 similarity,^36^ the MSforIDv2 similarity,^37^ the spectral entropy,^38^ the Scribe fragmentation accuracy,^39^* the Pearson correlation of fragment ion intensities,* the Spearman correlation of fragment ion intensities,* the Manhattan distance, the Euclidean distance, the Chebyshev distance, the Bray-Curtis distance, the Canberra distance, and the Ruzicka similarity. Versions of the spectrum similarity measures marked with “*” were also included when only considering peak matches that cover the five most intense fragment ions in the library spectrum.

ANN-SoLo supports two classification models for semi-supervised learning: a linear SVM, which closely mimics the original Percolator model,^6,33^ and a non-linear random forest classifier. Prior to rescoring, the PSM features, as described above, are preprocessed by removing the mean and scaling to unit variance (standardization), removing zero-variance features, and removing features that are highly correlated with other features (Pearson correlation above 0.95). Next, hyperparameter tuning is performed in a three-fold cross-validation set-up to evaluate different weights for the positive and negative classes (0.1, 1, or 10) for both types of classifiers, and evaluate different tree depths for the random forest. The classifier is subsequently trained using mokapot for multiple semi-supervised learning iterations to distinguish between true positive target PSMs, and false positive target PSMs and decoy PSMs, respectively. Finally, q-values are assigned to the PSMs based on their ranking by the learned mokapot scores.

As ANN-SoLo implements a cascade search strategy^32^ to maximize the identification of both unmodified and modified peptides,^29^ semi-supervised rescoring is performed separately for each stage of the cascade search. In the first stage of the cascade search, a standard search is performed using a small precursor mass tolerance to identify unmodified peptides. After mokapot rescoring, PSMs below the FDR threshold are extracted, while the spectra that remain unidentified are submitted to the second stage of the cascade search to identify modified peptides by performing an open search using a wide precursor mass tolerance. In this case, an additional feature is provided for each PSM to indicate its modification group membership. Modification groups consist of PSMs with similar precursor mass differences, to enable the classification model to learn different patterns for distinct PTMs. Modification groups are formed by frequently occurring precursor delta masses. These are determined by detecting peaks within the histogram of PSM precursor mass differences, split by 1 Da intervals around each nominal mass value, and assigning each PSM to its closest peak (while making sure to not violate the precursor mass tolerance). Modification groups are required to consist of at least 100 PSMs, with the remaining PSMs assigned to a residual group. Next, after training the classification model, for the open search q-value calculation is performed within each modification group separately, akin to a subgroup FDR strategy.^40^ Finally, PSMs from the first and second stage of the cascade search are combined to form the full spectrum identification results.

### 2.2 Data

A previously generated dataset measured from the HEK293 human cell line^23^ was used to evaluate the performance of ANN-SoLo rescoring. As per Chick et al. [23], the HEK293 cells were first lysed, trypsinized, and separated into 24 fractions, after which high-resolution and high-mass accuracy MS/MS spectra were obtained on an LTQ Orbitrap Elite mass spectrometer. For full details on the sample preparation and acquisition see the original publication by Chick et al. [23]. As input to the various search engines, previously generated Mascot Generic Format (MGF) files corresponding to this dataset were retrieved from the PRoteomics IDEntifications (PRIDE) database^41^ (project PXD009861).

For spectral library searching, the MassIVE-KB peptide library (version 2018/06/15)^42^ was used to search the data. This is a repository-wide human higher-energy collisional dissociation spectral library derived from over 30 TB of human MS/MS proteomics data. The original spectral library contained 2148 752 MS/MS spectra, from which duplicates were removed using SpectraST (version 5.0)^43^ by retaining only the best replicate spectrum for each individual peptide ion, resulting in a spectral library containing 2 113 413 spectra. Next, decoy spectra were added in a 1:1 ratio using the shuffle-and-reposition method,^44^ resulting in a final spectral library containing 4 226 826 spectra.

For sequence database searching, the human UniProt/SwissProt database was used (UP000005640, version 2022_06 containing 20 371 reviewed protein sequences),^45^ with decoys added using Philosopher.^46^

All MS/MS data, spectral libraries, sequence databases, and identification results have been deposited to the ProteomeXchange Consortium^47^ via the MassIVE partner repository with the dataset identifier RMSV000000197.3.

### 2.3 Search settings

#### 2.3.1 ANN-SoLo

ANN-SoLo version 0.4.0 was used to produce all search results. Spectra were preprocessed by removing the precursor ion peak and noise peaks with an intensity below 1% of the base peak intensity. If applicable, spectra were further restricted to their 50 most intense peaks. Spectra that contained fewer than 10 peaks remaining or with a mass range less than 250*m/z* after peak removal were discarded. Finally, peak intensities were rank transformed to deemphasize overly dominant peaks. A precursor mass tolerance of 10 ppm was used for standard searching, and a precursor mass tolerance of 10 ppm followed by 500 Da was used for cascade open searching. In both cases a fragment mass tolerance of 0.05 Da was used. FDR filtering was performed by the integrated functionality within ANN-SoLo, as described previously.

#### 2.3.2 SpectraST

We compared the performance of ANN-SoLo against the popular SpectraST spectral library search engine^43^ (version 5.0 as part of the Trans-Proteomic Pipeline version 5.1.0^48^). Spectra were preprocessed to have a minimum mass range of 250 Da, and only the 50 most intense peaks were retained. Peak-to-peak matching and rank-based scoring were used to evaluate spectrum matches. The precursor mass tolerance was 0.02 Da and the fragment mass tolerance was 0.02 Da. FDR filtering was performed externally by ranking the PSMs by the SpectraST dot product and using the plus-one correction^49^ during FDR calculation.

#### 2.3.3 COSS

We also compared the performance of ANN-SoLo against the recently developed COSS spectral library search engine (version 2.0).^50^ The cosine similarity was used as the scoring function, and the precursor mass tolerance and fragment mass tolerance were 10 ppm and 0.05 Da, respectively. The remaining settings were kept at their default values. FDR filtering was performed externally by ranking the PSMs by the COSS score (i.e. cosine similarity) and using the plus-one correction^49^ during FDR calculation.

#### 2.3.4 MSFragger

Finally, we compared the performance of ANN-SoLo against the state-of-the-art OMS sequence database search engine MSFragger (version 3.5).^26^

Because MSFragger is a sequence database search engine, whereas the previous tools are spectral library search engines, settings for MSFragger necessarily deviate slightly. MSFragger used the UniProt/SwissProt sequence database, rather than the MassIVE-KB spectral library, as described above. MSFragger was configured to consider tryptic peptides with up to 2 missed cleavages, and cysteine carbamidomethylation was specified as a static modification. A precursor mass window of 10 ppm was used for standard searching, and a precursor mass window of 500 Da was used for open searching. In both cases a fragment mass tolerance of 0.05 Da was used. For MSFragger results without PSM rescoring, FDR filtering was performed externally by ranking the PSMs by the MSFragger score (i.e. hyperscore) and using the plus-one correction^49^ during FDR calculation.

MSFragger results can be easily rescored by Percolator^6^ or PeptideProphet^51^ using the FragPipe user interface. Here we used Percolator (version 3.05.0)^6^ for PSM rescoring and q-values reported by Percolator were used. Note that FragPipe also has an option to calculate additional features for rescoring using the MS-Booster mode.^18^ However, because MS-Booster can only be used for standard searching, not for open searching, for consistency this functionality was not enabled.

### 2.4 Code availability

The ANN-SoLo spectral library search engine is available as an open-source Python command-line tool. It uses Pyteomics (version 4.5.5)^52^ to read MS/MS spectra; spectrum_utils (version 0.3.5)^53^ for spectrum preprocessing; Faiss (version 1.5.3)^54^ for efficient similarity searching; Scikit-Learn (version 1.0)^55^ and mokapot (version 0.8.0)^33^ for semi-supervised learning; and NumPy (version 1.20.3),^56^ SciPy (version 1.7.3),^57^ Numba (version 0.54.1),^58^ and Pandas (version 1.3.4)^59^ for scientific computing. Matplotlib (version 3.5.1)^60^ and Seaborn (version 0.11.2)^61^ were used for data visualization. Data analysis was performed using Jupyter notebooks.^62^

ANN-SoLo is available as open source under the permissive Apache 2.0 license on GitHub at https://github.com/bittremieux/ANN-SoLo. Analysis notebooks to reproduce the presented results are available on GitHub at https://github.com/issararab/ML-Results-Analysis-Notebooks.

## 3 Results

### 3.1 PSM rescoring boosts identification results of standard searching

When comparing the performance of ANN-SoLo to alternative spectral library search engines, SpectraST and COSS, ANN-SoLo is able to identify the largest number of MS/MS spectra, outperforming COSS and SpectraST on our benchmark dataset by 20% to 30%, respectively (figure 1a-b). Rescoring of the ANN-SoLo results further increases its performance advantage compared to SpectraST and COSS, albeit to a modest extent, by identifying 2.3% additional spectra at 1% FDR. Note that search engines explicitly need to support PSM rescoring by providing informative features that describe the quality of a spectrum match to be used for machine learning. Unfortunately, SpectraST currently does not provide such functionality. Additionally, although an integration between COSS and Percolator was recently described,^12^ this rescoring failed because COSS produced empty intermediate files to be used by Percolator. As such, only direct identification results could be reported for SpectraST and COSS, without PSM rescoring.

**Figure 1:**
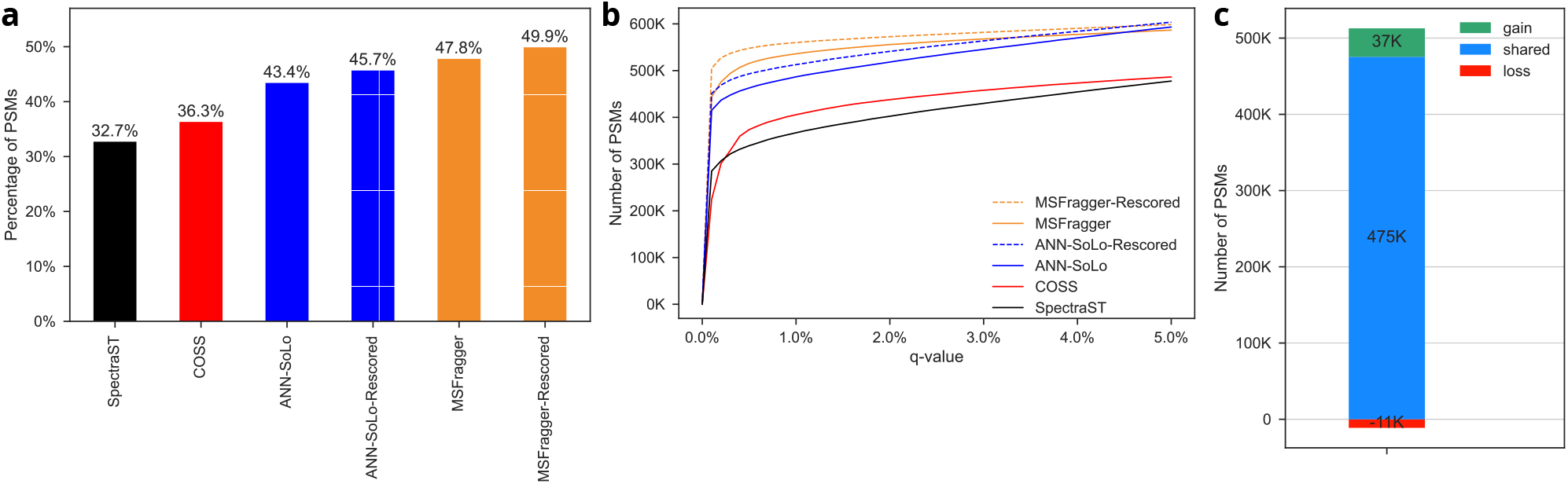
Spectrum identification performance for the HEK293 benchmark dataset, consisting of 1.1 million MS/MS spectra, using standard searching. **(a)** Fraction of identified spectra at 1% FDR. ANN-SoLo and MSFragger outperform SpectraST and COSS, with PSM rescoring providing an additional performance advantage for the former two search engines. **(b)** Spectrum identification performance at different FDR thresholds, visualized as a q-value plot.^63^ **(c)** Change in identified spectra by ANN-SoLo at 1% FDR after PSM rescoring.

We also compared the performance of ANN-SoLo to the popular MSFragger sequence database search engine (figure 1a-b). On average, MSFragger is able to identify slightly more spectra than ANN-SoLo. However, ANN-SoLo searched against the MassIVE-KB spectral library, which contains MS/MS spectra for less than 1 million unique peptidoforms, whereas MSFragger used a sequence database for the full human proteome, from which almost 3 million potential tryptic peptides could be generated. As such, the search space available for both tools differed, with 74% of peptides that were detected by MSFragger but not by ANN-SoLo were missing from the MassIVE-KB spectral library, and thus could never be found by ANN-SoLo. Irrespective of the difference in search space, when evaluating the PSM rescoring performance of ANN-SoLo and MSFragger, both tools were able to increase the number of confident PSMs by approximately 2% of all spectra at 1% FDR, corresponding to an increase of 26 074 and 23 545 in the number of identified spectra for ANN-SoLo and MSFragger, respectively.

When looking at how PSM rescoring changes the spectra that could be identified, we observe that the vast majority of PSMs were retained, in addition to the loss of a small number of PSMs and a larger gain of other PSMs (figure 1c). Furthermore, the majority of PSMs that were lost had higher q-values prior to rescoring, and thus were of lower confidence, whereas the q-values of the PSMs that were gained are uniformly distributed, indicating an unbiased gain in PSMs (Supplementary figure S1). Consequently, PSM rescoring not only increased the number of identified spectra, but also dropped lower-quality spectrum annotations to achieve more confident identification results.

### 3.2 PSM rescoring is a promising strategy for open searching

Besides their strong performance during standard searching, one of the advantages of ANN-SoLo and MSFragger is that both tools are able to perform open searches very efficiently. In contrast, because SpectraST and COSS do not have an efficient open searching mode and do not support PSM rescoring, they were omitted from the comparison. Although PSM rescoring is an increasingly ubiquitous strategy to improve the quality of standard searches, it has not been extensively used to post-process identification results from open searching yet. Therefore, we set out to explore the performance of PSM rescoring for open searching.

As expected, open searching was able to identify significantly more spectra than standard searching (figure 2a). Notably, whereas during standard searching MSFragger slightly outperformed ANN-SoLo, both tools obtain near-identical performance during open searching. Additionally, both tools benefited from PSM rescoring, and this benefit was stronger than during standard searching, with a PSM increase of 5.9% and 8.7% at 1% FDR for ANN-SoLo and MSFragger, respectively (figure 2a-b).

**Figure 2:**
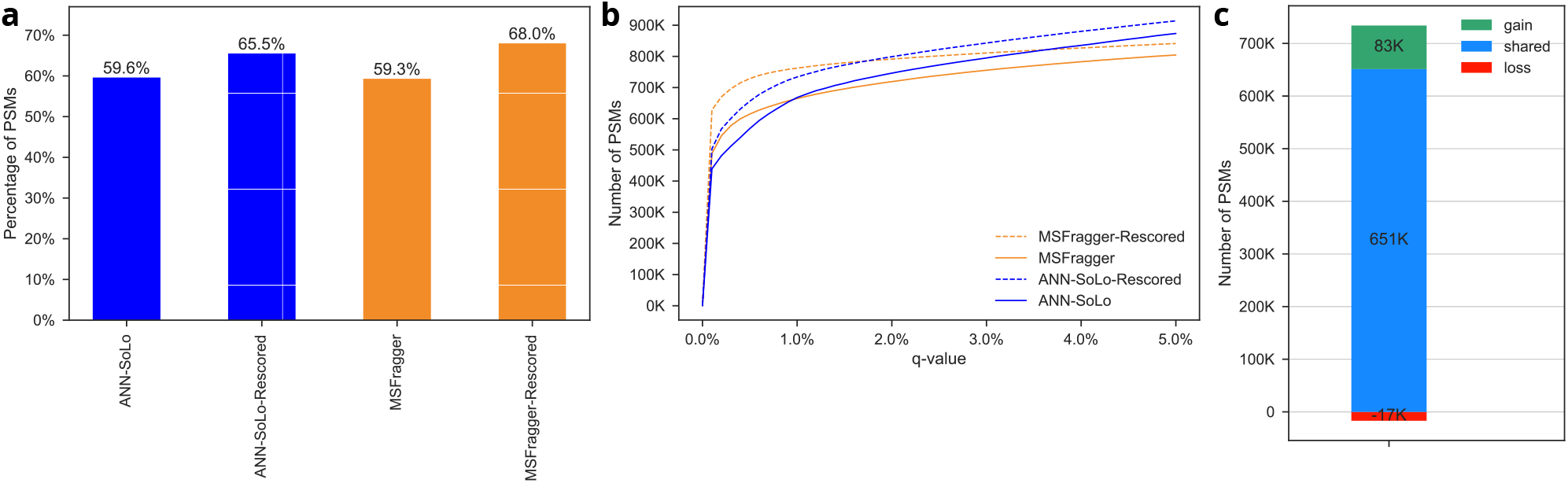
Spectrum identification performance for the HEK293 benchmark dataset, consisting of 1.1 million MS/MS spectra, using open searching. **(a)** Fraction of identified spectra at 1% FDR. PSM rescoring significantly increases the number of identified spectra for ANN-SoLo and MSFragger. **(b)** Spectrum identification performance at different FDR thresholds, visualized using a q-value plot.^63^ **(c)** Change in identified spectra by ANN-SoLo at 1% FDR after PSM rescoring.

Similarly to standard searching, PSM rescoring removed a small number of lower-quality PSMs and added a larger number of newly identified PSMs (figure 2c). When looking at the precursor mass differences of the newly included PSMs, we can derive that they correspond to several biologically relevant PTMs and chemical modifications introduced during sample processing,^64^ including phosphorylation, (di)oxidation, formylation, amidation, loss of ammonia, aminoethylbenzenesulfonylation, replacement of 2 protons by iron, etc. (Supplementary figure S2).

### 3.3 Effect of machine learning features for PSM rescoring of different search modes

A key aspect of the PSM rescoring performance is that informative features should be provided to the machine learning model so that it can successfully learn to discriminate between correct and incorrect spectrum identifications. To explore the relevance of various features, we investigated their importances in the SVM used by ANN-SoLo for PSM rescoring. The weights learned by a linear SVM for each feature can be used as a surrogates for the importances of those features (barring interference from correlated features).

During standard searching, several spectrum similarity measures receive large feature importances (figure 3a). For example, the cosine similarity, a ubiquitous spectrum similarity score,^65^ and which was used as the default score in ANN-SoLo previously;^29^ the Kendall-Tau correspondence between the intensity ranks of matched fragment ions and the hypergeometric probability of obtaining more than the observed number of matched fragment ions by random chance, which are used in the Pepitome spectral library search engine;^35^ and the MSforIDv1 score,^36^ which was introduced for matching of small molecule MS/MS spectra are among the top-ranked features. Although several features capture similar PSM aspects and show strong correlations with each other (Supplementary figure S3), nevertheless they each still contribute relevant information to the classifier.

**Figure 3:**
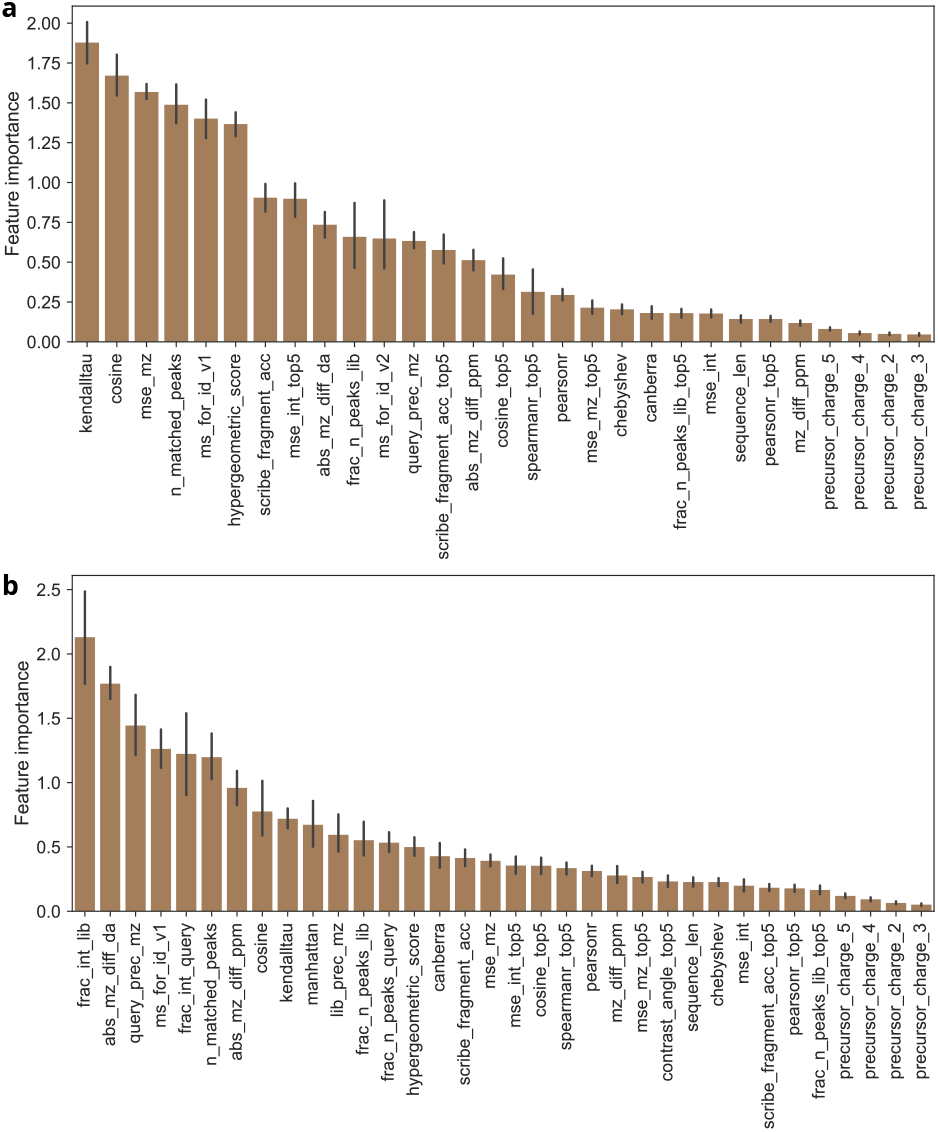
Absolute feature importances derived from the linear SVM used for PSM rescoring. **(a)** Feature importances for 29 features used during standard searching after redundant and uninformative feature removal. **(b)** Feature importances for 34 features used during open searching after redundant and uninformative feature removal.

Interestingly, other features were most important for PSM rescoring during open searching (figure 3b), including the absolute precursor *m/z* difference, the query precursor m/z, and the fraction of explained intensity in the query and library spectra. Whereas all candidate PSMs adhered to the 10 ppm precursor mass tolerance during standard searching, during the open search an essentially unrestricted precursor mass tolerance was used. Consequently, features capturing precursor m/z information received low feature importances during the standard search but high feature importances during the open search, as in the latter case the classifier could harness these features to learn patterns corresponding to specific PTMs (Supplementary figure S2). We can also observe that the feature correlations among each other are lower during open searching than during standard searching (Supplementary figure S4), indicating the importance of employing diverse and information-rich features for PSM rescoring.

## 4 Conclusions

Here we have demonstrated how PSM rescoring integrated in the ANN-SoLo search engine boosts the sensitivity and accuracy of open modification spectral library searching. Originally introduced by Percolator,^6^ PSM rescoring has grown into an increasingly popular approach to maximize the number of confident PSMs that can be obtained. Although PSM rescoring has so far predominantly been used to post-process identification results from standard sequence database searching, it was also recently introduced for post-processing of standard spectral library searching.^12^ We further expand on this by using PSM rescoring for both standard searching and open searching. Interestingly, our results show that different types of features capturing various PSM properties are relevant for different search modes. Notably, to rescore open searching results, grouping information to distinguish between PTMs, as derived from the precursor mass differences between the observed and theoretical masses, was provided to the classifier and a sub-group FDR approach was used to assign the rescored PSM q-values. Our evaluations show that such relevant features allowed the classifier to learn different PTM patterns and improve the identification of modified peptides from open searching.

PSM rescoring was fully integrated in ANN-SoLo using the mokapot library,^33^ which provides a flexible reimplementation of the Percolator algorithm in Python. Whereas other tools that have recently incorporated PSM rescoring have typically done so by using Percolator as an external post-processing tool, directly integrating rescoring in the ANN-SoLo codebase afforded us several advantages. First, because PSM rescoring is an integral part of ANN-SoLo, no external tools are used and there is no need for intermediate files to communicate between different tools. Consequently, PSM rescoring is fully transparent for the end user, which significantly improves the robustness and user-friendliness of this novel functionality. Second, being able to access the internals of the PSM rescoring functionality provided additional flexibility to include advanced functionalities. For example, PSM rescoring in ANN-SoLo can trivially switch between the standard linear SVM, as used by Percolator,^6^ or a random forest (Supplementary figure S5), which can learn a non-linear decision function and which has previously been shown to have beneficial performance during PSM rescoring.^8,33^ Additionally, other machine learning models, such as neural networks,^9^ could effortlessly be integrated as well if they support the popular Scikit-Learn application programming interface (API).^55^ Furthermore, being able to directly manipulate the PSM rescoring functionality allowed us to get deeper insights into the workings of the machine learning model, for example, to perform a detailed analysis of the feature importances. In contrast, when Percolator is used as an external, black box model, such information is much less accessible.

PSM rescoring is available via various tools, such as Percolator,^6^ PeptideProphet,^7^ Scavager,^8^ mokapot,^33^ and several custom approaches, and it is compatible with multiple proteomics search engines.^10,11^ A recent trend is to use predicted fragment ion intensities, often coming from deep learning tools,^66^ to compute a novel category of features to be used by Percolator or alternative PSM rescoring tools.^13–20^ Interestingly, similar features can be directly computed from spectral library searching results without the need for additional MS/MS spectrum prediction by complex deep learning tools. During spectral library searching, spectrum identifications are obtained by matching against real library MS/MS spectra, rather than against theoretical, simulated MS/MS spectra, providing detailed fragment intensity information virtually for free. As such, PSM rescoring is a natural approach to improve the sensitivity of spectral library searching.

PSM rescoring is now available in the opensource ANN-SoLo spectral library search engine to provide best-in-class spectrum identification performance at minimal additional complexity to the user. More information and detailed user instructions can be found at https://github.com/bittremieux/ANN-SoLo.

## Supporting information

**Supplementary Figure S1:**
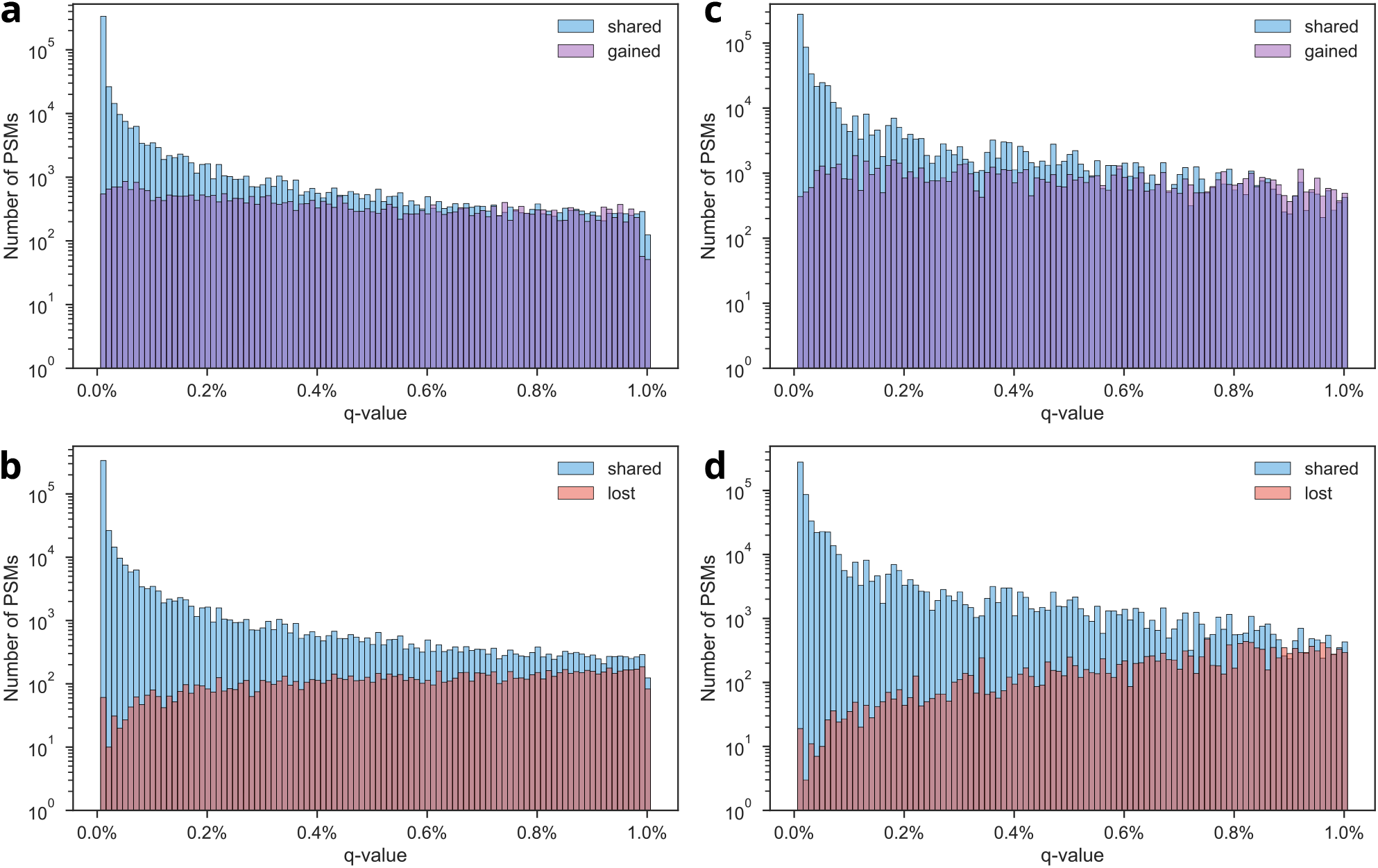
Q-value distributions for PSMs gained and lost during PSM rescoring, compared to the retained PSMs. **(a)** PSMs gained during standard searching. **(b)** PSMs lost during standard searching. **(c)** PSMs gained during open searching. **(d)**PSMs lost during open searching.

**Supplementary Figure S2:**
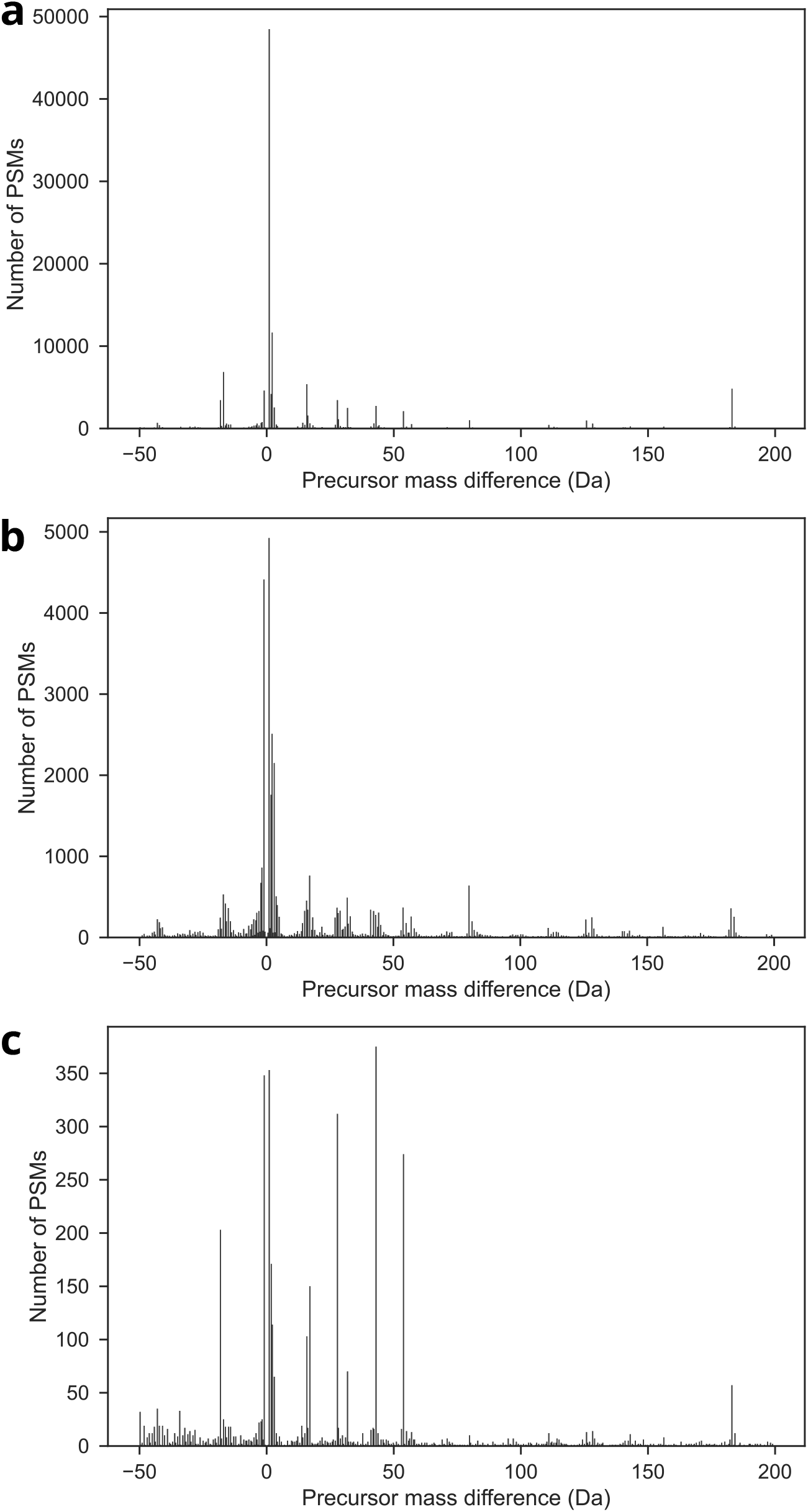
Histogram of non-zero precursor mass differences from PSMs at 1% FDR obtained from open searching. **(a)** Precursor mass differences for PSMs retained after PSM rescoring. **(b)** Precursor mass differences for PSMs gained after PSM rescoring. **(c)** Precursor mass differences for PSMs lost after PSM rescoring.

**Supplementary Figure S3:**
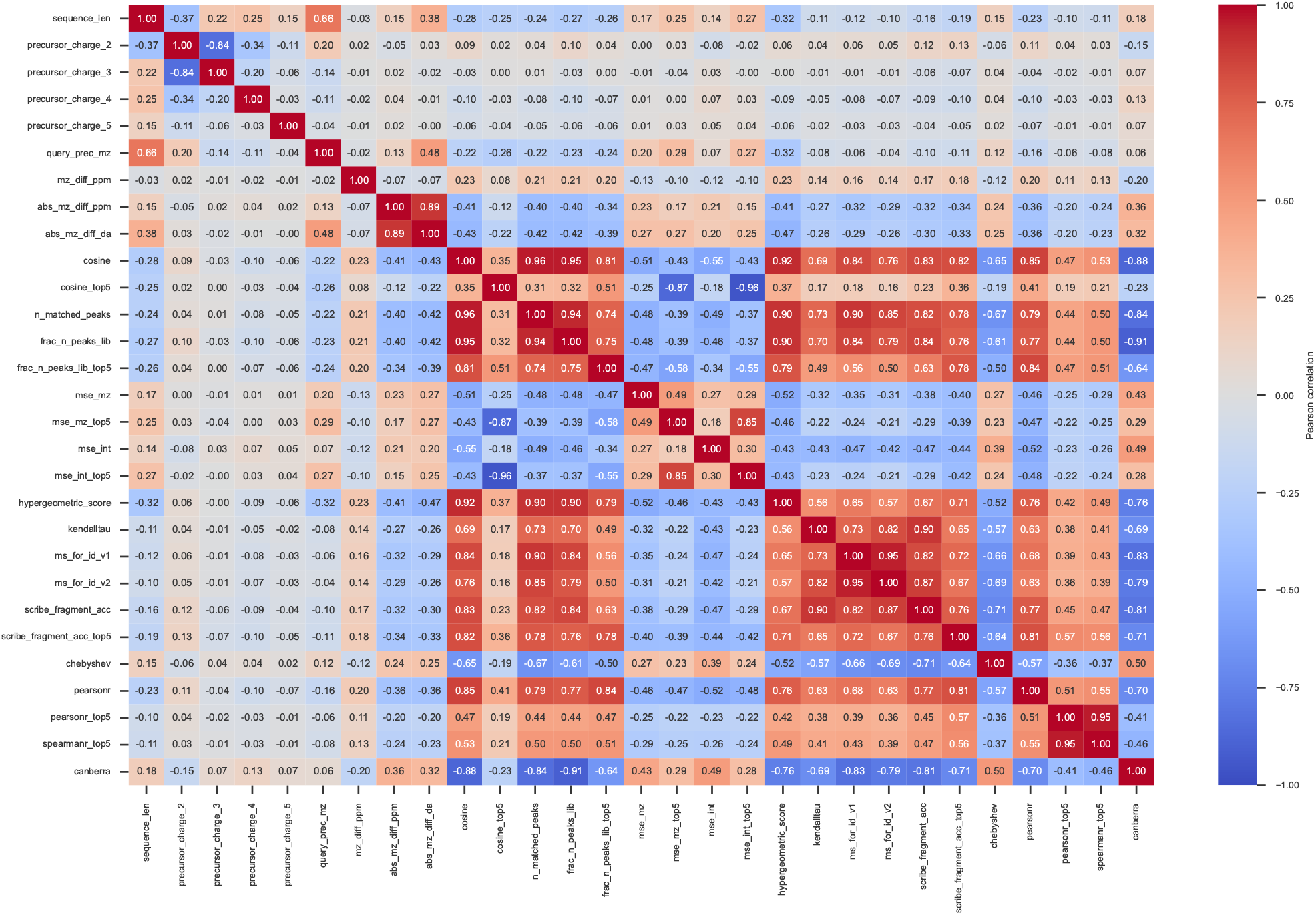
Correlations among 29 features used during standard searching after redundant and uninformative feature removal.

**Supplementary Figure S4:**
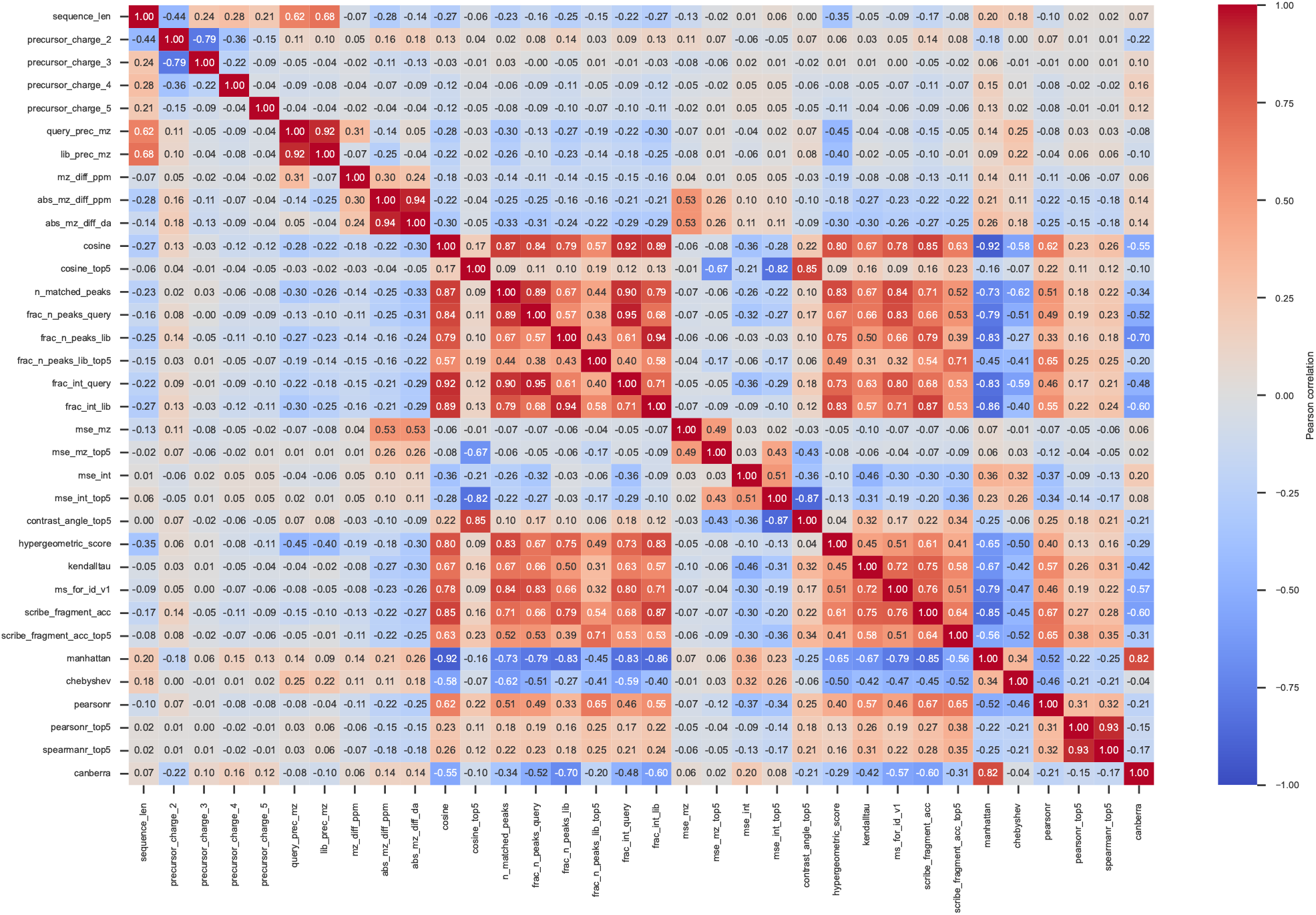
Correlations among 34 features used during open searching after redundant and uninformative feature removal.

**Supplementary Figure S5:**
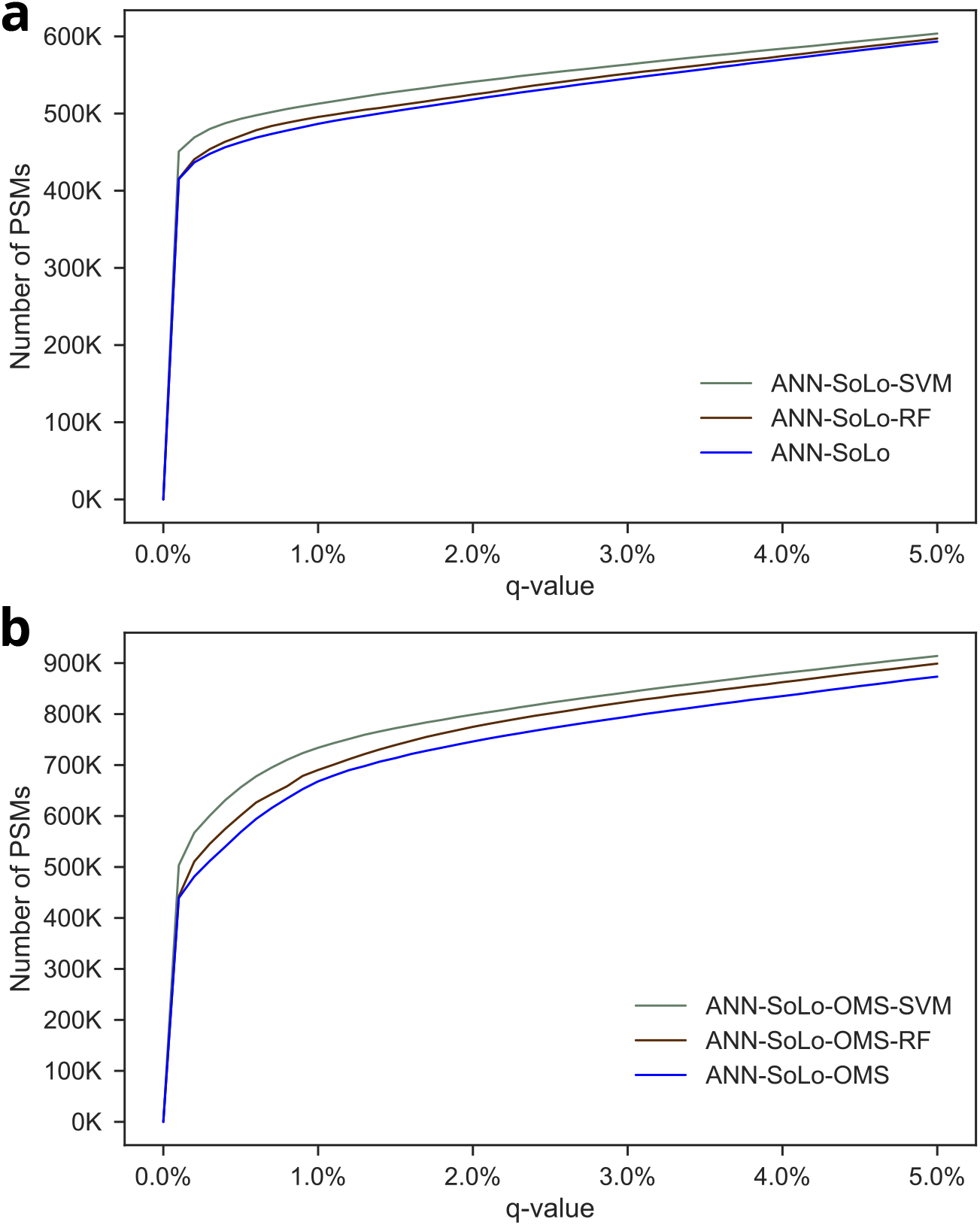
Spectrum identification performance for the HEK293 benchmark dataset, consisting of 1.1 million MS/MS spectra, using standard searching. Two different classifiers, a linear SVM and a random forest, were used for PSM rescoring, with the SVM consistently achieving the best performance. **(a)** Standard searching. **(b)** Open searching.

## Notes

https://github.com/bittremieux/ANN-SoLo

https://massive.ucsd.edu/ProteoSAFe/reanalysis.jsp?task=d64e84b87b35469e9eb2faa4563c3b3f

https://www.ebi.ac.uk/pride/archive/projects/PXD009861

### Competing Interest Statement

The authors have declared no competing interest.

